# The Conserved C-terminal End Segment of Troponin T Is A New Tropomyosin-Binding Site Modulating the Kinetics of Cardiac Muscle

**DOI:** 10.1101/2023.02.20.529301

**Authors:** Tianxin Cao, Han-Zhong Feng, J.-P. Jin

## Abstract

Evolved from duplication of a troponin I (TnI)-like ancestor gene, troponin T (TnT) and TnI are two subunits of the troponin complex that regulates the contraction and relaxation of striated muscles. Proteolytic deletion of the evolutionarily added N-terminal variable region of TnT in adaptation to acute myocardial stress restores a conformation like that of the C-terminal end segment of TnI, a tropomyosin (Tm)-binding and inhibitory structure, with an effect on reducing the contractile velocity of cardiac muscle to prolong ejection time and sustain the stroke volume. To investigate the underlying mechanism of this adaptive conditional function of TnT that is known to have two Tm-binding sites, our study localized a conformationally modulated new Tm-binding site in the highly conserved 14 amino acid C-terminal end segment of TnT. Localized surface plasmon resonance data showed that a hypertrophic cardiomyopathy mutation R278C within the C-terminal end segment of TnT alters the tropomyosin binding with direct responses to physiological concentrations of Ca^2+^. The functions are retained in the form of free peptide with an inhibitory regulatory effect on the contractile and relaxation kinetics of skinned cardiac muscle. In addition to revealing the underlying mechanisms of cTnT-ND adaptation and cardiac TnT C-terminal myopathic mutations, the new findings provide novel insights into the structure-function relationship of TnT in the kinetics of striated muscle contraction and relaxation with broad physiological and pathophysiological implications.

## INTRODUCTION

The contraction and relaxation of vertebrate striated muscles (i.e., skeletal and cardiac muscles) are regulated by Ca^2+^ via the troponin (Tn) complex in sarcomeric thin filaments (Gordon, Homsher et al. 2000, Tobacman 2021). Troponin consists of three subunits: troponin C (TnC, the calcium binding subunit), troponin I (TnI, the inhibitory subunit), and troponin T (TnT, the tropomyosin (Tm)-binding subunit). During the activation of contraction, cytosolic Ca^2+^ increases and binds to TnC to induce a series of conformational changes in Tn and Tm and open the thin filament to activate thick filament myosin ATPase and cross bridge cycling. Directly interacting with TnC, TnI and Tm, TnT plays a central role in the regulation of striated muscle contraction and relaxation by anchoring the Tn complex to Tm and transmitting the Ca^2+^ induced conformational signals (Perry 1998, Wei and Jin 2016).

The central and C-terminal regions of TnT are highly conserved in structure and contain two binding sites for Tm (Mak and Smillie 1981, Jin and Chong 2010). The N-terminal region of TnT has highly variable sequences among muscle type-specific isoforms and across species and does not contain known protein binding sites (Perry 1998, Wei and Jin 2016). Myocardial ischemia-reperfusion induces a restrictive truncation of the N-terminal variable region (residue 1-71) of cardiac TnT preserving all conserved structures (Zhang, Biesiadecki et al. 2006). The N-terminal truncated cardiac TnT (cTnT-ND) remains in the myofilament with functional effects. Transgenic mouse hearts expressing the cTnT-ND to replace the endogenous intact cTnT demonstrated reduced systolic velocity and extended left ventricular rapid-ejection time to sustain the stroke volume when afterload increases (Feng, Biesiadecki et al. 2008). This finding demonstrate that cTnT-ND is a posttranslational mechanism of compensatory adaption by gaining an inhibitory function to modulate the contractile kinetics of cardiac muscle.

TnT and TnI evolved from duplication of a TnI-like ancestor gene (Chong and Jin 2009). An intriguing observation is that deletion of the evolutionarily added N-terminal variable region of TnT restores a sub-molecular conformation recognized by a monoclonal antibody (mAb) TnI-1 raised against the C-terminal end segment of TnI (Jin, Yang et al. 2001, Chong and Jin 2009). This finding indicates that a TnI-like ancestral conformation is resumed in cTnT-ND after removing the evolutionary added N-terminal segment that is a known conformational tuning structure (Wang and Jin 1998, Biesiadecki, Chong et al. 2007). The mAb TnI-1 recognized C-terminal end segment of TnI is a Ca^2+^-regulated allosteric Tm-binding structure in the troponin complex (Zhang, Akhter et al. 2011). The C-terminal end 27 amino acids segment of cardiac TnI (cTnI-C27) binds Tm in the form of free peptide with functional effect on reducing the Ca^2+^ sensitivity of activated cardiac muscle strips (Wong, Feng et al. 2019, Hornos, Feng et al. 2021).

To investigate the cTnI-C27-like inhibitory structure resumed in cTnT-ND as an underlying mechanism for the compensatory effect of cTnT-ND on reducing ventricular contractile velocity to elongate ejection time and sustain stroke volume in myocardial energetic crisis (Feng, Biesiadecki et al. 2008), the present study identified a conformationally modulated new Tm-binding site in the highly conserved 14 amino acid C-terminal end segment of TnT. Myopathic mutations can alter the property of this new Tm-binding site of cardiac TnT with direct responses to physiological levels of Ca^2+^. The functions are retained in the form of isolated 14 amino acids C-terminal peptide of cardiac TnT with inhibitory regulatory effects on the contractile and relaxation kinetics of skinned cardiac muscle. In addition to revealing the underlying mechanisms of cTnT-ND adaptation and cardiac TnT C-terminal myopathic mutations, the new findings provide novel insights into the structure-function relationship of TnT in the kinetics of striated muscle contraction and relaxation with broad physiological and pathophysiological implications.

## MATERIALS AND METHODS

### Protein constructs and synthetic peptides

Fig. 1A summarizes the TnT protein constructs used in our study with the functional sites and mAb epitopes annotated. cDNAs encoding intact and fragments of TnT are constructed in pAED4 or pET17b plasmid vectors for expression in *E. coli*. The following constructs are produced: Intact adult mouse cardiac TnT, N-terminal truncated mouse cardiac TnT (McTnT-ND72, engineered by polymerase chain reaction, PCR, as described previously (Biesiadecki, Chong et al. 2007), Recombinant pET17b plasmids encoding wild type and mutant human mini-cTnTs that contain the exons-15-17 encoded C-terminal 62 amino acids were purchased from Gene Universal Inc. (Newark, DE).

**Figure 1.**
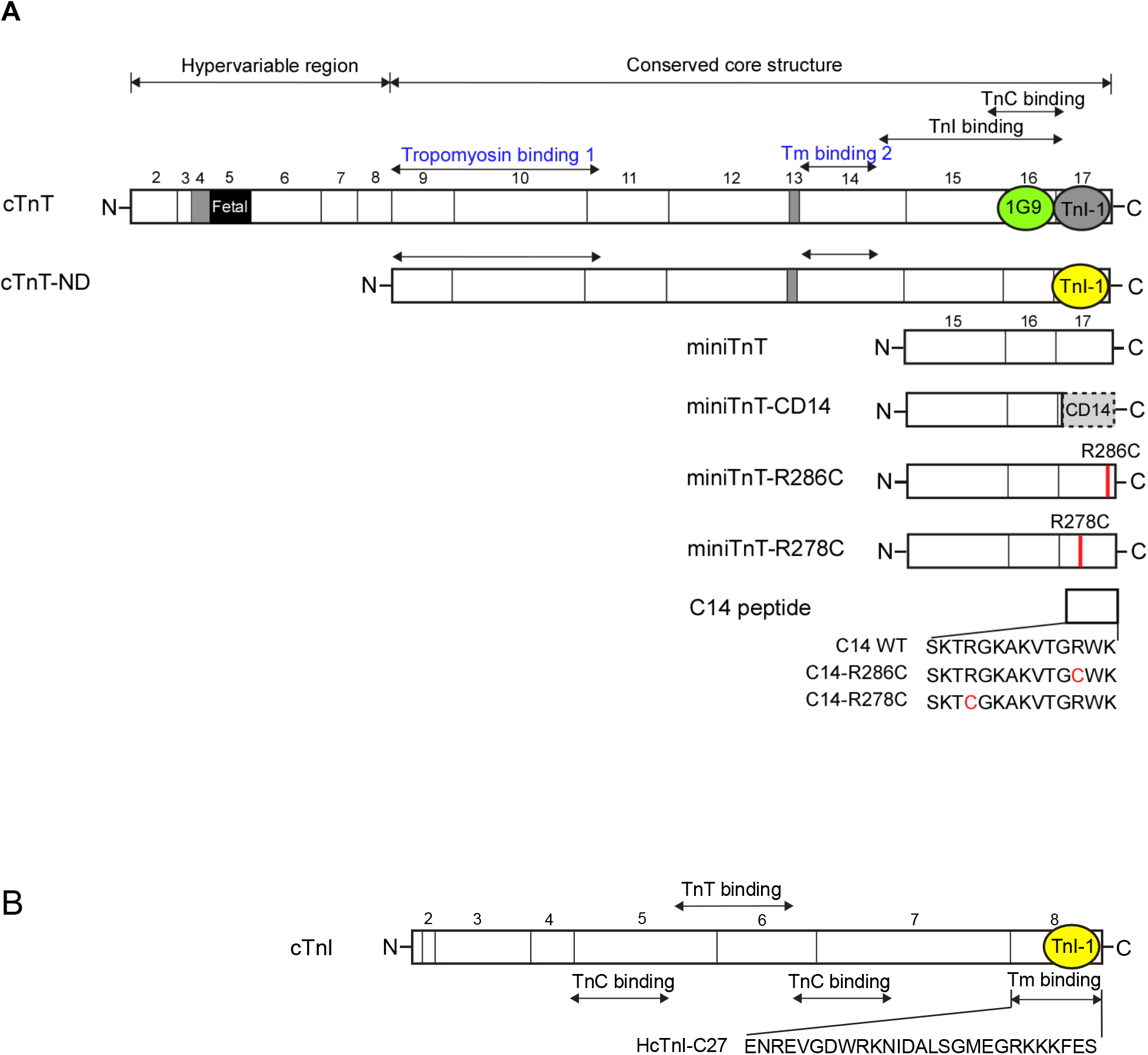
TnT constructs used in the present study. (A) Primary structural maps of mini-TnT constructs analyzed are shown in alignment with intact cardiac TnT and cTnT-ND. The previously identified Tm, TnI and TnC binding sites and the location of epitopes recognized by mAbs 1G9 and TnI-1 are indicated on the linear structural map. The sequences of WT and mutant HcTnT-C14 peptides encoded by the last exon (exon 17) are also shown. (B) The amino acid sequence of the C-terminal end segment of TnI C27 shows no apparent similarity to that of the C-terminal end segment of TnT, indicating the critical role of folded conformation in the functionality of proteins.

The cDNA constructs were expressed by transformation of BL21(DE3)pLysS *E. coli* cells. 37°C cultures in 2x TY medium containing 100 mg/L ampicillin and 12.5 mg/L chloramphenicol were induced with 0.4 mM isopropyl-1-thiol-β-D-glactoside at mid-log phase for 3 additional hours for protein expression. The bacterial cells were harvested by centrifugation at 4°C, re-suspended in a lysis buffer containing 2.5 mM EDTA and 50 mM tris-HCl, pH 8.0, and lysed using French Press.

The purification of intact cardiac TnT and McTnT-ND72 was described previously (Biesiadecki, Chong et al. 2007). The Mini-TnTs expressed were fractionated using a G75 gel filtration column in 6 M urea, 0.5 M KCl, 0.1 mM EDTA, 6 mM β-mercaptoethanol, 10 mM imidazole-HCl, pH 7.0. Fractions containing mini-TnT were dialyzed against 0.1mM EDTA and lyophilized. All purification steps were carried out at 4°C.

Free peptides HcTnI-C27 (Fig. 1B) (Wong, Feng et al. 2019), HcTnT-C14 WT, HcTnT-C14 R278C and HcTnT-C14 R286C (Fig. 1A) were synthesized by using a commercial service (Peptide 2.0 Inc., Chantilly, VA) at >95% purity.

### Site-specific anti-TnT and anti-TnI monoclonal antibodies

Using standard hybridoma technology, a mouse mAb TnI-1 was generated by immunization using purified chicken fast skeletal muscle TnI as described previously (Jin, Yang et al. 2001). mAb TnI-1 specifically recognize a conserved epitope at the C-terminal end segment (Fig. 1A) of all three muscle type TnI isoforms in all vertebrate species tested (Jin, Yang et al. 2001).

A mouse mAb 1G9 was generated by immunization using purified human cardiac TnT. Previous characterization showed it is against an epitope in the C-terminal T2 region (Chong and Jin 2009) (Fig. 1A).

mAb TnI-1 (IgG1) was purified from mouse hybridoma ascites fluid using Protein G chromatography (Millipore Sigma P7700). TnI-1 IgG eluted with 50 mM glycine-HCl, pH 2.7, was neutralized immediately with 1M Tris-HCl, pH 8, and dialyzed again phosphate-buffered saline (PBS) for experimental use.

### SDS-polyacrylamide gel electrophoresis (SDS-PAGE)

SDS-PAGE using various gel formulas was carried out as described previously (Jin 1995). Protein samples were prepared in sample buffer containing 2% SDS and 1% β-mercaptoethanol. 14% Laemmli gels with an acrylamide:bisacrylamide ratio of 29:1 were used to resolve intact TnT and TnT fragments. 15% Tris-Tricine small pore SDS-gels (Schagger and von Jagow 1987) with acrylamide:bisacrylamide ratio of 20:1 were used to resolve mini-TnTs and small peptides. The resolved gels were stained with Coomassie Blue R-250.

### Microtiter plate mAb epitope analysis

Enzyme-linked immunosorbent assay (ELISA) was performed to examine the conformational changes in the C-terminal end segment of TnT using mAbs TnI-1 and 1G9 as epitope affinity probes (Rasmussen and Jin 2022). Purified TnI and TnT proteins were dissolved and coated at 2.5 μg/mL in Buffer A (0.1 M KCl, 3 mM MgCl_2_, 20 mM Imidazole, pH 7.0, 10mM EGTA) containing either 0.003 mM Ca^2+^ (pCa 9) or 10.1 mM Ca^2+^ (pCa 4) as calculated using UC Davis Ca-Mg-ATP-EGTA v1.0 computer program. Proteins were coated 100 μL/well at 4°C overnight. After removing unbound proteins by washing with Buffer A containing 0.05% Tween-20 (Buffer T) and blocking the plates with 150 μL/well Buffer B (Buffer T containing 1% BSA) of pCa 9 or 4 at room temperature for 30 min, the immobilized TnT was incubated with 100 μL/well serial dilutions of TnI-1 mAb in Buffer T containing 0.1% BSA (Buffer D) of pCa 9 or 4 at room temperature for 2 hrs. After washing with Buffer T, the plates were incubated with 100 μL/well horseradish peroxidase-conjugated anti-mouse immunoglobulin second antibody (Sigma, St. Louis, MO) in Buffer D of pCa 9 or 4 at room temperature for 1 h. Unbound secondary antibody was removed by three washes as above. The binding of mAb probe to the TnT constructs was detected by H_2_O_2_/2,2’-azinobis-(3-ethylbenzthiazolinesulfonic acid) substrate reaction. The color development in each assay well was monitored at a series of time points using an automated microplate reader (BioTek, Synergy-HTX). A_420nm_ values in the linear course of color development were used to plot titration curves and the antibody dilution required for 50% maximum binding was quantify for the binding affinity of mAb TnI-1 to its specific epitope in the TnT constructs. All experiments were done in triplet wells.

### Microtiter plate protein binding assay

ELISA solid-phase protein–binding experiments were performed to assess the binding affinity of TnT C-terminal end segment to rabbit a-tropomyosin. As described previously (Biesiadecki and Jin 2011), purified TnT proteins were coated on 96-well microtiter plates at 5 μg/mL, 100 μL/well in Buffer A at 4°C overnight as described above. The plates were washed with Buffer T and blocked with Buffer B at room temperature for 30 min. Serial dilutions of rabbit α-Tm were added in Buffer D to incubate at room temperature for 2 hrs. The binding of Tm was then detected by anti-α-Tm mAb CH1 (Lin, Chou et al. 1985) at room temperature for 1 hr. After washing with Buffer T as above to remove unbound mAb CH1, HRP-conjugated anti-mouse immunoglobulin secondary antibody was added for incubation of 45 min used to detect binding of the primary antibody. After final washes as above, H_2_O_2_/2,2’-azinobis-(3-ethylbenzthiazolinesulfonic acid) substrate was added for color development at room temperature. Absorbance at 420 nm was monitored using an automated microplate reader at 5 min intervals for 30 min. Except for the substrate development, pCa of the reactions was controlled at 9 and 4 as described in the mAb epitope analysis above. The assays were done in triplicate wells. TnT-Tm binding curves were constructed using data collected at a time point prior to the end of the linear phase of color development.

### Localized surface plasmon resonance (LSPR) epitope and binding analysis

Using an Open SPR^™^ instrument (Nicoya Lifesciences, Canada), LSPR was employed to study binding of the C-terminal end segment of TnT to Tm. Purified Tm was covalently immobilized on a carboxyl sensor chip of following the chip activation via 1-ethyl-3-(3-dimethylaminopropyl) carbodiimide (EDC) and N-hydroxysuccinimide (NHS) as described previously (Rasmussen and Jin 2022). A blocking solution (Nicoya Lifesciences, Canada) was injected to block any remaining activated esters on the chip. Purified mini-TnT and peptides were diluted at 20μM in the running buffer (10 mM piperazine-N,N’-bis(2-ethanesulfonic acid), pH 7, 100 mM KCl, 3 mM MgCl_2_, 10 mM EGTA) and injected at a flow rate of 20 μL/min over 300s to measure the binding affinity Ka for immobilized Tm, following by injecting running buffer at 20 μL/min over 300s to measure the dissociation rate Kd. The data were used to further determine the binding kinetic rate constant K_D_ (Kd/Ka). The chip was regenerated with 0.5 M KCl between runs. The effect of Ca^2+^ was analyzed by controlling the solutions at pCa 9 (0.003 mM Ca^2+^) or pCa 4 (10.01 mM Ca^2+^) as described above.

### Contractility of skinned cardiac muscle sections

Studies of mouse cardiac muscle contractility were performed using protocols approved by the Institutional Animal Use and Care Committee (ACC) of University of Illinois at Chicago.

Permeabilized mouse cardiac muscle strips were prepared using a skinned cryosection method described previously (Feng and Jin 2020). Left ventricular papillary muscles were dissected and rapid frozen in optimal cutting temperature (O.C.T.) compound by submerging in liquid nitrogen. The frozen papillary muscle was sectioned longitudinally into 35 μm thick and 120-150 μm wide strips using a cryostat. The muscle strips collected were washed with a relaxing buffer (40 mM BES, 10 mM EGTA, 6.86 mM MgCl_2_, 5.96 mM ATP, 1 mM DTT, 33 mM creatine phosphate, 200U/mL creatine kinase, 3.28 mM K-propionate, pH 7.0, plus protease inhibitor cocktail) and stored in a 35 mm dish at −20°C in relaxing buffer containing 50% glycerol until the use in force-pCa studies.

Operated on a thermal-controlled metal stage at 0°C, the cryo-sectioned cardiac muscle strips selected with cardiomyocytes clearly organized along the long axis were mounted between two aluminum T-clips and transferred in relaxation buffer to an 8-chamber stage (802D, Aurora Scientific) thermo-controlled at 6-8°C mounted on an inverted microscope. The muscle preparation was connected to a force transducer (403A, Aurora Scientific) and a length controller (322C, Aurora Scientific). The buffer was then switched to a skinning solution (relaxation buffer plus 1% Triton X-100) for 20 min to further permeabilize the muscle strips. After a wash with relaxation buffer without Triton X-100, the permeabilized cardiac muscle strip was placed in pCa 9.0 buffer made by mixing the relaxing buffer (pCa 10.0) with a pCa 4.0 activation buffer (40 mM BES, 10 mM EGTA, 6.64 mM MgCl_2_, 6.23 mM ATP, 1 mM DTT, 10 mM CaCl_2_, 33 mM creatine phosphate, 200U/mL creatine kinase, 2.09 mM K-propionate, pH 7.0, plus protease inhibitor cocktail) and sarcomere length was measured by direct imaging using a digital camera attached to the microscope and adjusted to 2.0 μm and 2.3 μm. Calcium activated force was measured at pCa 6.5, 6.3, 6.0, 5.8, 5.5, 5.0, and 4.5 at 15°C as described previously (Feng and Jin 2020). The series of pCa buffers were made by mixing the relaxation buffer and activation buffer with the free [Ca^2+^] calculated using Fabiato’s program (Fabiato and Fabiato 1979). HcTnT-C14 peptide was then added to 20 μM and the force-pCa measurements were repeated. The force-pCa curves were plotted and fitted using Hill exponential equation.

### Phylogenetic analysis

Amino acid sequences were analyzed using DNAStar MegAlign Pro software (Lasergene Inc, Madison, WI). Sequence alignment was generated by Clustal W method and phylogenetic tree of representative animal species was constructed using maximum-likelihood method algorithm.

### Data analysis

Statistical analysis of LSPR results were compared for Ka, Kd, and KD by using Student’s t test. All values are presented as mean ± SD.

## RESULTS

### Expression and purification of a set of mini-TnT proteins

To investigate the epitope conformation and biochemical interaction between Tm and C-terminus cTnT, mini-TnT encoding exons 15-17 of HcTnT (Fig. 1A) was expressed in *E.coli* and purified (Fig. 2) to remove the N-terminal conformational effect and the two previously identified high affinity Tm binding sites (Jin and Chong 2010). To further narrow down the epitope site, a mini-TnT with last 14 amino acid deletion (miniTnT-CD14) was constructed (Fig. 1 and 2). C-terminal last 14 amino acid is a product of a G > A mutation in the splice donor sequence of intron 15, resulting in the truncation that causes familial cardiac hypertrophy (Mukherjea, Tong et al. 1999). Other hypertrophic cardiomyopathy mutations R286C (Richard, Charron et al. 2003) and R278C (Morimoto, Nakaura et al. 1999) with various degrees of severity within the mini-TnT region were also included in this study. Due to the high-level expression of the set of mini-TnT proteins, they were all formed inclusion bodies after French press lysis (Fig. 2). Only gel filtration chromatography was needed to purify the protein.

**Figure 2.**
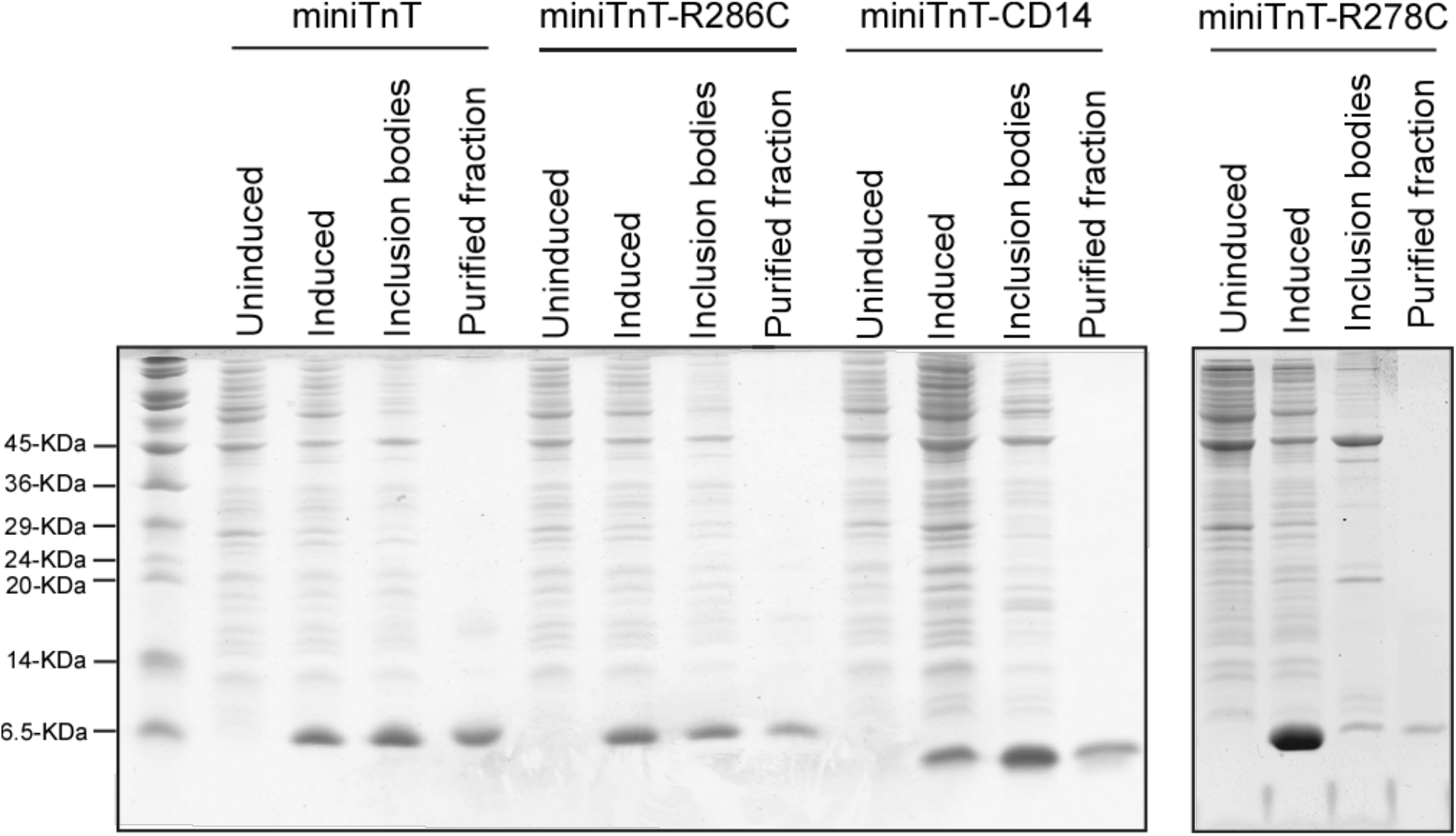
Expression and purification of WT, C-terminal truncated and mutant mini-TnT constructs. The Tris-Tricine small pore 15% SDS-gels show the bacterial expressions and purified preparations of mini-TnT-WT, mini-TnT-CD14, mini-TnT-CD2, mini-TnT-R278C and mini-TnT-R286C.

### cTnT-ND restores a TnI-like epitope structure in the C-terminal end segment of cTnT

The TnI-1 mAb (Jin, Yang et al. 2001) previously generated against an epitope in the C-terminal end segment of TnI has no cross-reaction to intact cardiac TnT but shows a high affinity binding to N-terminal truncated cardiac TnT (Fig. 3). This conditional structural similarity between TnT and TnI is consistent with the common ancestry of TnI and TnT genes and indicates the presence of an evolutionarily repressed TnI-like molecular conformation in TnT (Chong and Jin 2009).

**Figure 3.**
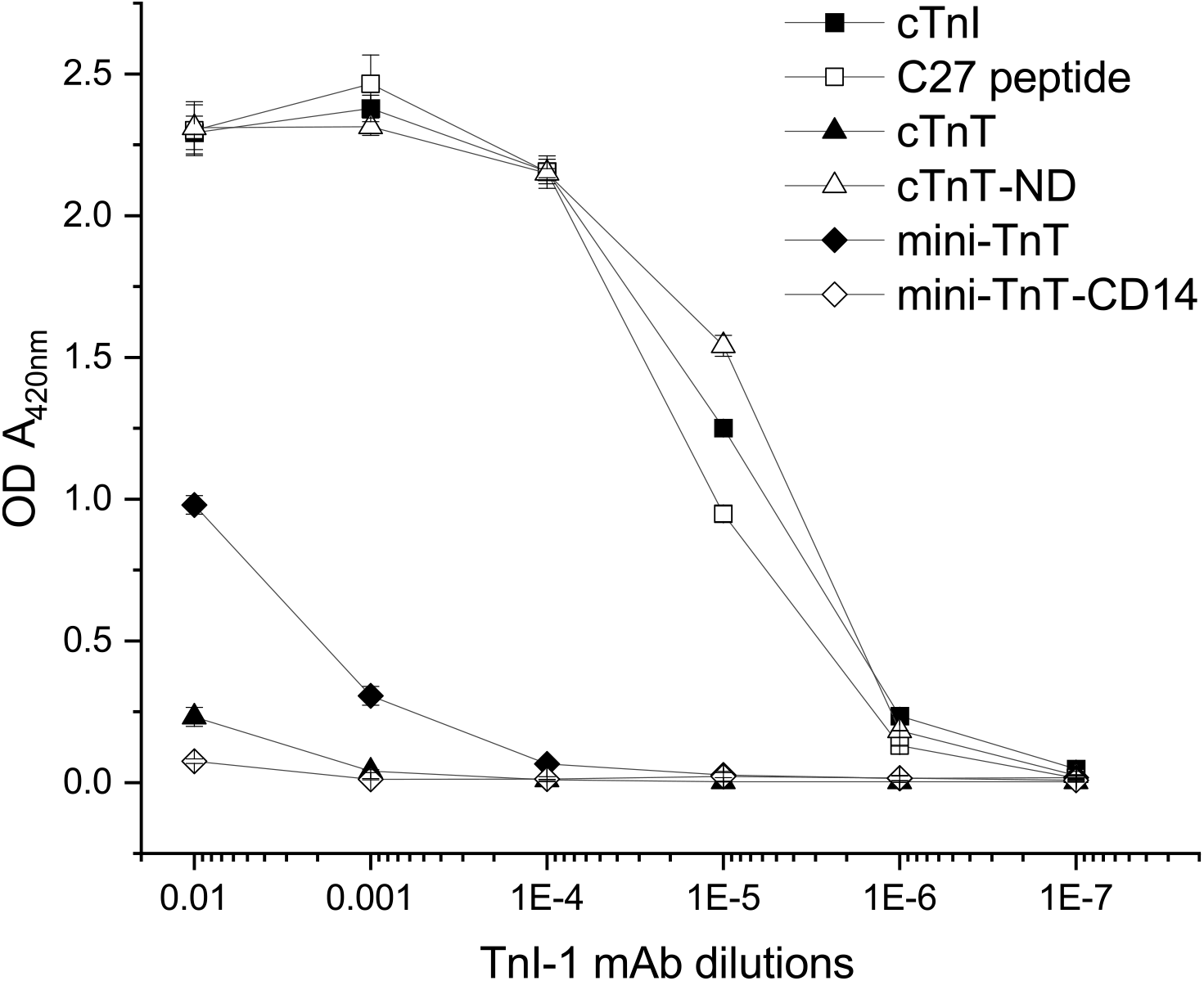
Identification of a conditional mAb TnI-1 epitope in the C-terminal end segment of TnT. The ELISA titration curves of mAb TnI-1 against microplate-immobilized cardiac TnT (cTnT) and mini-TnT constructs together with intact mouse cardiac TnI (cTnI) and human cardiac TnI C-terminal peptide (C27) controls showed that mAb TnI-1 raised against the C-terminal end segment of TnI has only no significant cross-reaction to intact cardiac TnT whereas cTnT-ND becomes strongly recognized by mAb TnI-1 with a high affinities nearly identical to that for TnI. This resumed TnI C terminus-like epitope conformation is readily detectable in mini-TnT but completely abolished by deletion of the C-terminal 14 amino acids, localizing the conditional TnI C terminus-like epitope in the C-terminal end segment of TnT. Since the amino acid sequence of the C-terminal end segment of TnI has no apparent sequence similarity to that of the C-terminal end segment of TnT (Fig. 1B), the conditional TnI-1 epitope in cTnT-ND is a folded structurebased function.

This conditional TnI-1 epitope of TnT is preserved in the C-terminal domain of cardiac TnT as shown by the strong cross-reactions of mAb TnI-1 to cTnT-ND and mini-TnT (Fig. 3) whereas not present in intact cTnT. Deletion of the C-terminal 14 amino acids from mini-TnT results in complete loss of the TnI-1 epitope (Fig. 3), localizing this conditional TnI-like epitope structure in the C-terminal end segment of TnT.

The C-terminal end segment of vertebrate TnT is encoded by the last exon with highly conserved amino acid sequence suggesting the functional importance of this region. The phylogenetic tree of the last exon encoded C terminus segment from different isoforms of TnTs across species is shown in Fig. 4A. Muscle type specific isoforms of TnT were clustered separately. Fast skeletal muscle TnTs are further away from slow and cardiac TnTs, suggesting that fast type TnTs are most likely the prototype among three types of TnTs. Amino acid sequences alignment demonstrates that the last 14 amino acid sequences in all mammalian cTnT sequenced to date are identical (Fig. 4B), indicating a common functionality involved in in mammalian hearts.

**Figure 4.**
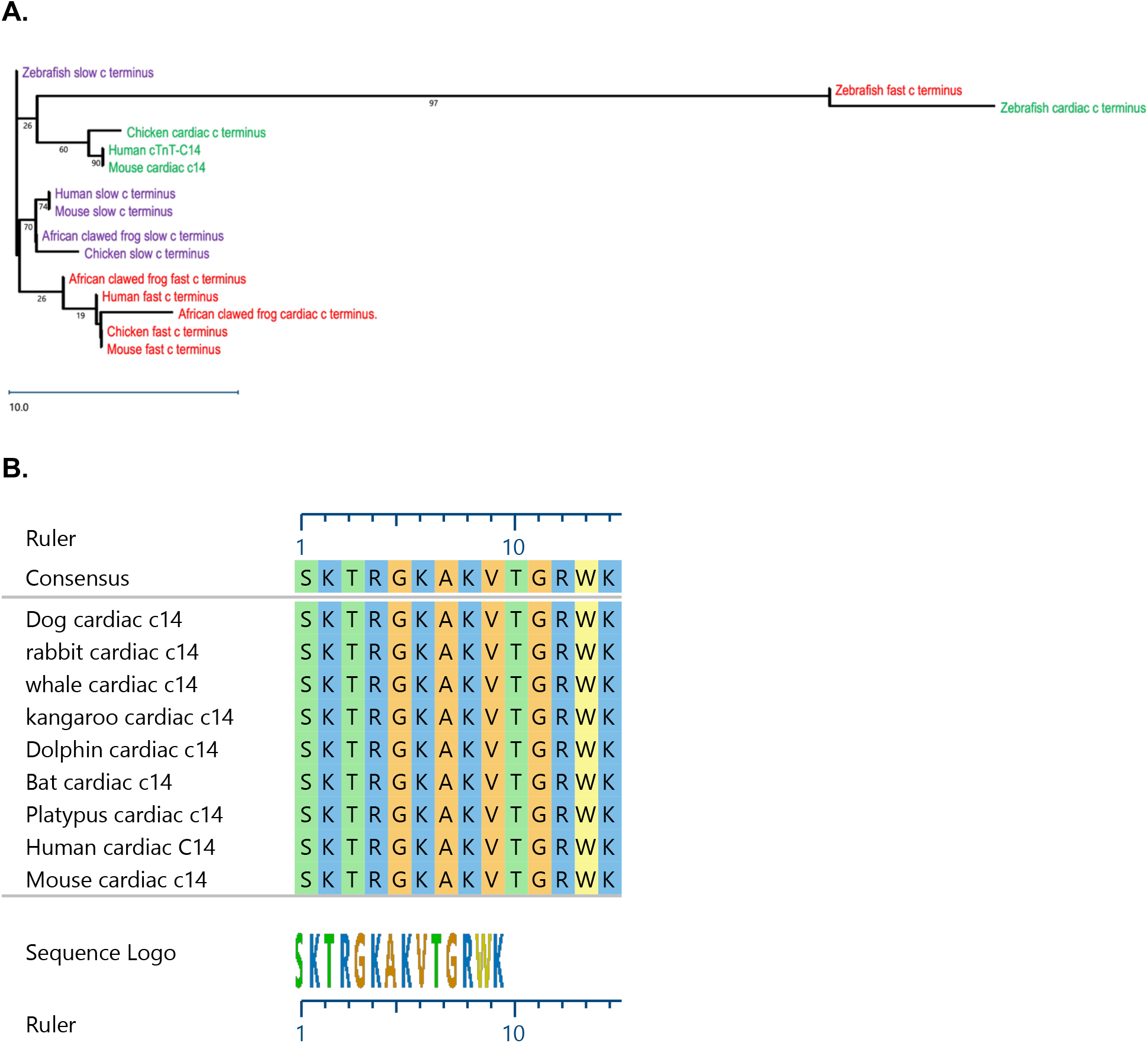
The C-terminal end segment of TnT is a conserved structure in muscle type isoforms across vertebrate phylum with identical sequence in mammalian cardiac TnTs. (A) The phylogenetic tree of amino acid sequences of the C-terminal end segment of cardiac, fast, and slow skeletal muscle TnTs from human, mouse, chicken, frog and fish was generated by alignment using DNAStar computer software. The evolutionary lineages demonstrate higher conservation of each of the muscle type isoforms among higher vertebrate species (birds and mammals) than that between isoforms. (B) Alignment of amino acid sequences of the C-terminal end segment of representative mammalian species. A striking phenomenon is that all mammalian cardiac TnT in the current database including marsupials have identical sequence in this segment encoded by the end exon. The accession numbers of the sequences shown are: Human, AAK92231.1; mouse, AAA85350.1; dog, NP_001003012.2; bovine, NP_777196.1; rabbit, XP_008266826.1; bat (XM_045817086), dolphin (XM_033846215.1), whale (XM_036872460), kangaroo (XM_013017336), platypus (XM_029069048).

### The 14 amino acids C-terminal end segment of TnT is a new tropomyosin binding site

We previously demonstrated that the C-terminal end 27 AA (C27) segment of TnI recognized by mAb TnI-1 binds Tm (Zhang, Akhter et al. 2011). The cTnI-C27 epitope structure was preserved in both isolated C27 peptide of TnI and in intact cTnI as detectable by mAb TnI-1 (Wong, Feng et al. 2019) and functions as an activated state myofilament Ca^2+^ desensitizer (Wong, Feng et al. 2019, Hornos, Feng et al. 2021). Therefore, we further tested whether the TnI-1 epitope structure resumed in the C-terminal end segment of cTnT also binds Tm.

We first performed Tm-binding assays with the set of mini-TnTs using ELISA at a muscle relaxation state (pCa 9.0). Fig. 5A showed that when the two known Tm-binding sites are removed, mini-TnT retains a saturable binding to Tm whereas mini-TnT-CD14 completely lost the binding activity. Our results demonstrate that there is a third Tm binding site exists within the last 14 AA of cTnT. Meanwhile, mini-TnTs with hypertrophic cardiomyopathy mutations R278C and R286C significantly reduced the Tn-binding affinity (Fig. 5A).

**Figure 5.**
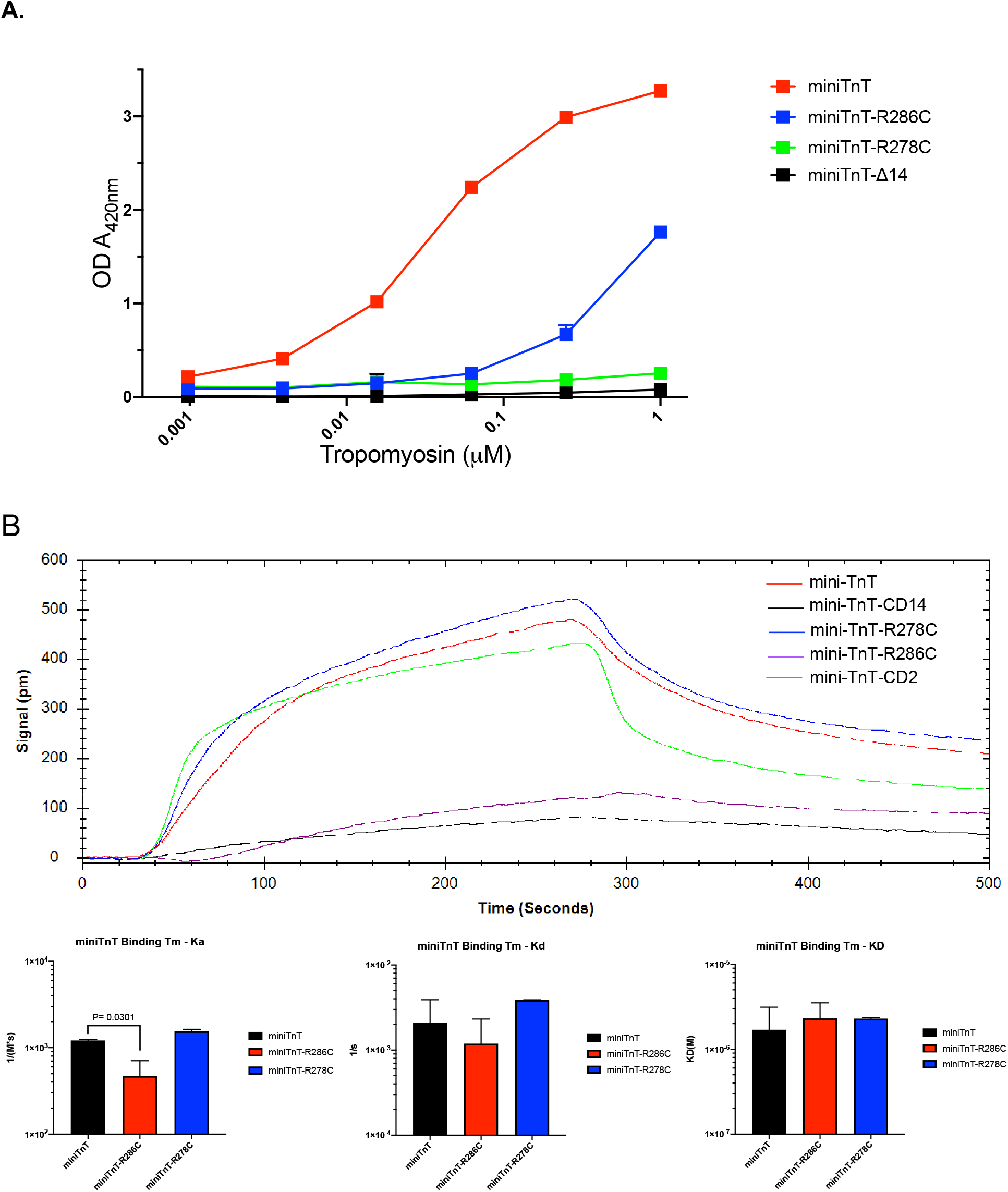
The C-terminal end segment of TnT is a new tropomyosin-binding site. ELISA protein-binding studies were performed at pCa 9 to examine the binding of Tm to mini-TnT constructs immobilized on microtitering plates. (A) The affinity titration curves detected that mini-TnT-WT shows a saturable strong binding to Tm, which is completely abolished by deletion of the C-terminal 14 amino acids in mini-TnT-CD14. Myopathic mutation R286C results in partial loss of the Tm-binding affinity while mutation R278C has a more drastic effect. The data demonstrate that the C-terminal end segment of 14 amino acids encoded by the last exon is a novel Tm-binding site in addition to the previously identified Sites 1 and 2 (Fig. 1). (B) LSPR characterization of the third Tm-binding site of TnT confirms the high Tm-binding affinity of mini-TnT-WT, which is abolished by deletion of the C-terminal end 14 amino acids in mini-TnT-CD14. Myopathic mutations showed differential altered Tm-binding affinities as compared to WT.

To determine the Tm binding kinetics of mini-TnT, LSPR was employed to detect the association rate (Ka), dissociation rate (Kd), and equilibrium dissociation constant (K_D_) in the relaxation state (pCa 9.0). Consistent with the results from the ELISA binding assay, deletion of the C-terminal 14 AA in mini-TnT abolished the binding with Tm, and the mini-TnT constructs with cardiomyopathic mutations showed significantly altered Tm-binding kinetics (Fig. 5B). Notably, while mini-TnT-R286C significantly lowered the maximum signal and association rate (Ka), R278C mutation increased the Tm-binding in LSPR (Fig. 5B).

### The third Tm binding site of cTnT becomes directly responsive to Ca^2+^ in myopathic mutants

Our previous study detected a Ca^2+^ regulated low but saturable Tm-binding by a tertiary troponin complex reconstituted with and TnC, C27-truncated cTnI and mini-TnT (Zhang, Akhter et al. 2011). To investigate whether the Ca^2+^ regulation retains in the cTnT-C14 segment, synthetic WT and mutant C14 peptides were studied. LSPR was used to demonstrate the binding between C14 and Tm at different Ca^2+^ concentrations. There was no significant difference in Tm-binding kinetics for WT cTNT-C14 peptide in the activation (pCa 4.0) and relaxation (pCa 9.0) states (Fig. 6). Interestingly, R278C mutation significantly increased the binding to Tm with much higher effect at pCa 9.0 than that at pCa 4.0.

**Figure 6.**
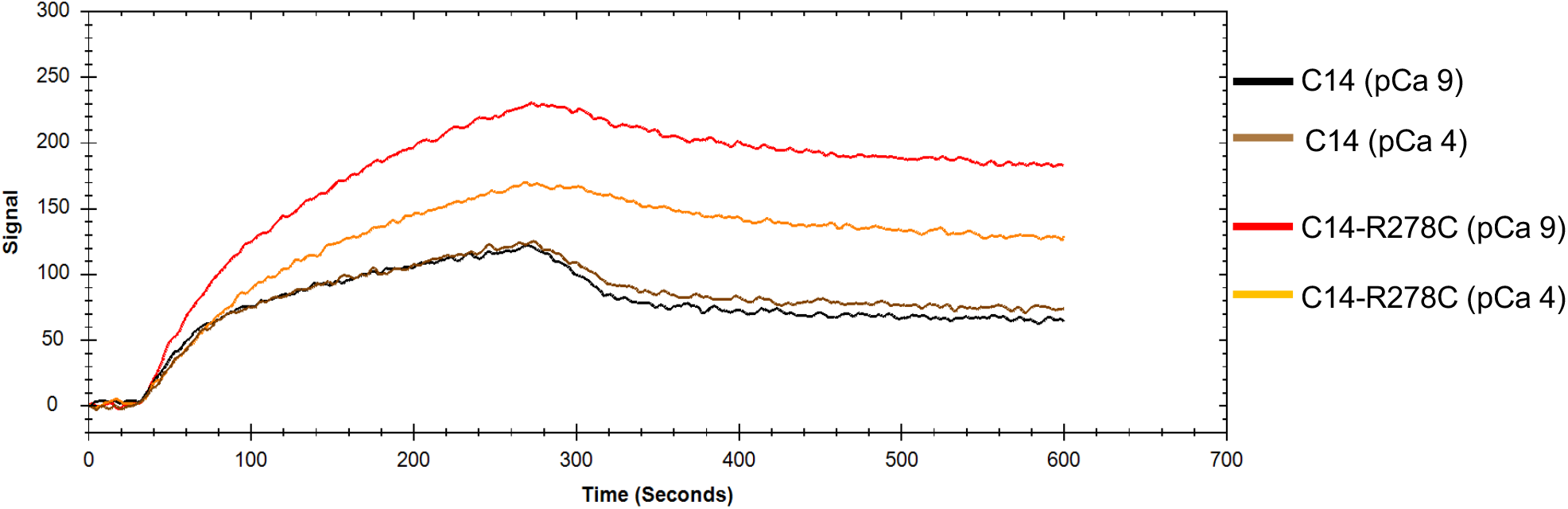
The third Tm-binding site of cardiac TnT became directly Ca^2+^-responsive in cardiomyopathic mutants. LSPR Tm-binding study showed that while there was no significant difference in Tm-binding kinetics for WT cTNT-C14 peptide in the activation (pCa 4.0) and relaxation (pCa 9.0) states (Fig. 6), R278C mutation significantly increased the binding to Tm with much higher effect at pCa 9.0 than that at pCa 4.0.

### Effect of C14 peptide on myofibril contractility

To further investigate the function of the C-terminal end segment in regulating cardiac muscle contractility, the effect of of cTnT-C14 peptide on the force-pCa relationship of cardiac muscle was studied using skinned mouse papillary muscle sections at sarcomere length 2.0 μm. The result in Fig. 7 showed that C14 peptide produced a right-shift of force-pCa curve corresponding to decreased pCa50 and calcium sensitivity, mimicking the effect of cTnI-C27 peptide on skinned muscle sections (Wong et al., 2019; Honors et al., 2021).

**Figure 7.**
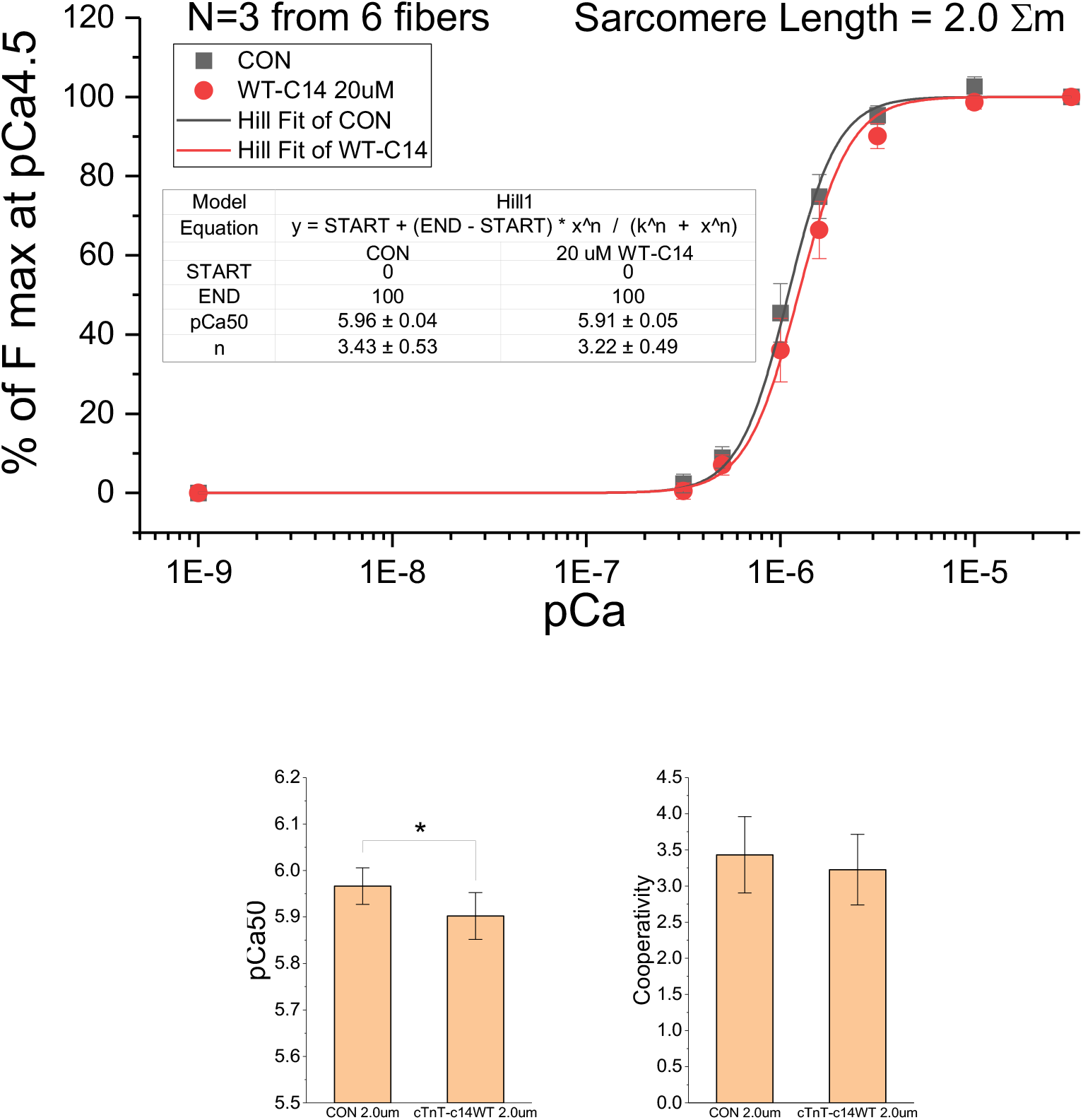
Functional effect of cTnT-C14 peptide on skinned cardiac muscles. Skinned mouse papillary muscle sections were used to measure the isometric force activation at a series of Ca^2+^ concentrations in the absence and presence of 20 μM WT cTnT-C14 peptide. The force-pCa curves show C14 peptide produced a right-shift, indicating decreased myofilament Cabsensitivity.

## DISCUSSION

### The C-terminal end segment of TnT is the third Tm-binding site

Both ELISA binding (Fig. 3) and LSPR (Fig. 5 and 6) studies demonstrated that there is a Tm-binding site presnt in the C-terminal end segment of mini-TnT. LSPR study further showed Tm-binding affinity with the C14 peptide of the last 14 AA of cTnT (Fig. 6). Previously, we have mapped and determined the two known strong Tm-binding sites of TnT using difference length of TnT fragments, localizing Site 1 in the middle region and Site 2 in the beginning of the T2 region (Fig. 1) (Jin and Chong 2010). Therefore, the C-terminal end Tm-binding site identified in the present study is a third Tm-binding site of TnT.

The C-terminal end segment of TnT is highly conserved in isoforms of TnT (Fig. 4A) with identical amino acid sequence among mammalian cardiac TnTs (Fig. 4B). This observation indicates an essential function of the C-terminal segment of TnT with a unique evolutionary selection for mammalian cardiac TnTs. Current crystal structure of troponin only resolved the core structure missing the C-terminus of TnT (Takeda, Yamashita et al. 2003). The lack of high-resolution structure of this segment reflects its flexible and allosteric nature. Supporting our findings, there were evidences that the C terminus of TnT may bind Tm near the Cys-190 site labeled with acrylodan probe (Morris and Lehrer 1984, Franklin, Baxley et al. 2012), which was weakened when Ca^2+^ binds TnC (Pearlstone and Smillie 1981).

### The C-terminal Tm-binding site is a conformationally modulated TnI-like inhibitory structure

Extended from the initial finding in a previous study (Chong and Jin, 2009), restrictive deletion of the N-terminal variable region (Fig. 1 A) made cardiac TnT positive to the anti-TnI mAb TnI-1 (Fig. 3). The TnI-1 epitope is originally located at the last 27 AA of TnI, functioning as myofilament Ca^2+^ desensitizer by binding to Tm (Wong, Feng et al. 2019). Removal of the evolutionarily additive hypervariable N-terminal domain of cardiac TnT as that occurs in the adaptation to acute myocardial ischemia or pressure overload resumed a TnI-like epitope structure that was not present in intact cTnT (Zhang, Biesiadecki et al. 2006, Chong and Jin 2009) with a functional effect on reducing the systolic velocity of the heart for compensatory increases of ventricular ejection time and stroke volume.

Mini-TnT fragment further shortened from cTnT-ND (Fig, 1A) had the C27-like epitope structure preserved as recognized by TnI-1 mAb in ELISA titrations (Fig. 3). The mini-TnT also retains the binding to Tm in both ELISA and LSPR binding assays (Fig. 5 and 6), which was mapped to the C-terminal end 14 AA with the mini-TnT-CD14 control (Fig. 5A and 6). Supporting our findings, Chalovich et al have studied the function of the C-terminal segment of cTnT (Gafurov, Fredricksen et al. 2004, Franklin, Baxley et al. 2012, Johnson, Angus et al. 2018). Their results showed that the C-terminal segment of TnT is a regulatory domain required for maintaining the inactive B-state for muscle relaxation and for limiting the active M-state at Ca^2+^-bound state.

### The C-terminal Tm-binding site with HCM mutation of TnT became directly responsive to _Ca_2+

While the third Tm-binding site in cTnT is not directly regulated by Ca^2+^ as shown in the LSPR study (Fig. 6), HCM mutation R278C significantly increased Tm-binding affinity of this segment and made it Ca^2+^ responsive. The binding activity is higher at pCa 9.0 than that at pCa 4.0 (Fig. 6). The stronger overall Tm-binding of cTnT-C14-R278C may indicate an over-inhibitory effect on myofilament activation which would reduce myocardial contractility.

In summary, after two Tm-binding sites have been identified in TnT (Perry, 1998; Jin and Chong, 2010), our study demonstrated the third Tm-binding site in the C-terminal end segment of TnT with TnI-like inhibitory function. This new Tm-binding site is conformationally modulated in cTnT-ND adaptation. The identification and functional characterization of this novel third conditional Tm-binding site of TnT laid a new foundation for our understanding of troponin structure-function relationship and the molecular mechanism of striated muscle contraction.

## Abbreviations used

BSA: bovine serum albumin
ELISA: enzyme-linked immunosorbant assay
LSPR: localized surface plasmon resonance
mAb: monoclonal antibody
SDS-PAGE: SDS-polyacrylamide gel electrophoresis
TBS: Tris-buffered saline
Tm: tropomyosin
Tn: troponin
TnC: troponin C
TnI: troponin I
TnT: troponin T

## ACKNOWLEDGMENTS

This study was supported in part by grants from the National Institutes of Health HL127691 and HL138007 (to J-PJ).

